# Overlap among spatial memories triggers repulsion of hippocampal representations

**DOI:** 10.1101/099226

**Authors:** Avi J. H. Chanales, Ashima Oza, Serra E. Favila, Brice A. Kuhl

## Abstract

Across the domains of spatial navigation and episodic memory, the hippocampus is thought to play a critical role in disambiguating (pattern separating) representations of overlapping events. However, the mechanisms underlying hippocampal pattern separation are not fully understood. Here, using a naturalistic route-learning paradigm and spatiotemporal pattern analysis of human fMRI data, we found that hippocampal representations of overlapping routes gradually diverged with learning to the point that they became less similar than representations of non-overlapping events. This representational ‘reversal’ of the objective route similarity (a) was selective to the hippocampus, (b) only occurred for the specific route segments that were shared across routes, and (c) was predicted by the degree to which individual hippocampal voxels were initially shared across route representations. These findings indicate that event overlap triggers a repulsion of hippocampal representations—a finding that provides critical mechanistic insight into how and why hippocampal representations become separated.

## INTRODUCTION

Distinct experiences often contain overlapping elements, creating the potential for memory interference. For example, a single location (e.g., a living room) may be the site of many different experiences and corresponding memories. The hippocampus is widely thought to play a critical role in coding overlapping events such that interference is minimized. Compelling evidence for this function comes from intracranial recordings in rodents during spatial navigation. For example, when rodents alternate between left-and right-hand turns in a T-maze, cells within the hippocampus differentially fire during the central stem (the overlapping path), according to whether the current route is a ‘right-turn’ or ‘left-turn’ route (Wood et al., 2000;Frank et al., 2000). Likewise, hippocampal place fields may completely remap with contextual changes in a rodent’s environment (Bostock et al., 1991;Colgin et al., 2008). In human studies of episodic memory, fMRI evidence indicates that visual stimuli that are shared across multiple event sequences are distinctly coded in the hippocampus according to the specific sequence to which they belong (Hsieh et al., 2014). While these studies and others have led to general agreement that the hippocampus forms distinct codes for overlapping experiences (Favila et al., 2016;Chadwick et al., 2010;Ginther et al., 2011;Agster et al., 2002;McKenzie et al., 2014;Brown et al., 2010;Kumaran and Maguire, 2006;Gilbert et al., 2001;Schlichting et al., 2015;Grieves et al., 2016;Kyle et al., 2015;LaRocque et al., 2013) the factors that trigger divergence of hippocampal representations are not fully understood.

The formation of distinct hippocampal representations is traditionally thought to be a result of sparse coding within the hippocampus (O’Reilly and McClelland, 1994;O’Reilly and Rudy, 2001;Marr, 1971;Leutgeb et al., 2007;McHugh et al., 2007;Bakker et al., 2008;Yassa and Stark, 2011;Treves and Rolls, 1994;McClelland and Goddard, 1996;GoodSmith et al., 2017). Although there are not enough neurons in the hippocampus to entirely avoid representational overlap, sparse coding ensures that similar experiences are less likely to share neural units, thereby resulting in orthogonalized representations. While this coding property of the hippocampus may play a critical role in reducing overlap during initial encoding, it is unlikely to provide a complete account of how hippocampal representations become distinct. In particular, overlap among hippocampal representations also changes with experience, suggesting learning-related factors that contribute to divergence. For example hippocampal remapping in rodents may emerge over the course of learning (Lever et al., 2002; Bostock et al., 1991), and even the sensitivity of stable hippocampal place fields can be tuned by experience (Mehta et al., 2000). Similarly, experience-dependent divergence of hippocampal activity patterns has been observed in human fMRI data (Favila et al., 2016;Schlichting et al., 2015;Schapiro et al., 2012;Kim et al., 2017). Computational models suggest that one factor that drives learning-related divergence of hippocampal representations is competition (Norman et al., 2006;Norman et al., 2007;Hulbert and Norman, 2014;Kim et al., 2017). That is, when activity patterns overlap–which may reflect residual overlap following initial orthogonalization–this overlap creates competition during learning that the hippocampus ‘solves’ by reducing similarity among representations. This perspective makes a critical prediction: that overlapping representations should systematically move apart from one another over the course of learning. Indeed, the representational distance between overlapping events should increase to a greater degree than the distance between non-overlapping events. This idea, which can be thought of as repulsion, is quite distinct from the idea of orthogonalization, because repulsion necessarily requires that an event’s representation is directly shaped by a similar (competing) event’s representation. Limited evidence from human fMRI studies hints at repulsion among overlapping hippocampal representation (Favila et al., 2016;Schlichting et al., 2015;Schapiro et al., 2012) but these observations come from episodic memory paradigms with static visual stimuli, which contrasts sharply with the spatial learning and navigation paradigms that have been used to study disambiguation of hippocampal activity patterns in rodents.

Here, we sought to bridge evidence from spatial learning paradigms in rodents and human episodic memory paradigms by testing, in a pair of human fMRI studies, whether overlap among spatial routes triggers an experience-dependent repulsion of hippocampal representations. Modeled after canonical rodent T-maze paradigms, we used a real-world route-learning paradigm that contained pairs of spatially-overlapping routes. However, in contrast to rodent T-maze paradigms, we also included pairs of non-overlapping routes, so that the similarity of overlapping route representations could be expressed relative to the similarity of non-overlapping route representations–a critical comparison for testing whether divergence preferentially occurs among overlapping routes. fMRI data were collected over the course of an extended learning session, allowing for representational similarity to be compared across time. Additionally, because our route stimuli were temporally dynamic, we used a novel spatiotemporal pattern analysis method wherein neural representations consisted of patterns of activity distributed across space (fMRI voxels) and time.

Our paradigm allowed us to test several critical predictions. First, if repulsion occurs, representations of overlapping events should diverge to a greater degree than non-overlapping events—that is, overlapping events should systematically move apart from each other. An unambiguous sign of repulsion is if overlapping event representations become less similar than non-overlapping event representations—what we will refer to as a ‘reversal effect’–as this outcome cannot be explained by orthogonalization of neural codes. Recently, we have shown at least one learning context in which a reversal effect is observed in the hippocampus (Favila et al., 2016), but it remains to be determined whether this seemingly paradoxical result is a general property of the hippocampus and whether it applies to the types of spatial learning paradigms commonly used in rodent studies. Second, to establish the critical point that event overlap itself triggers repulsion of hippocampal representations, it is essential to establish that repulsion only occurs for the segments of routes that actually overlap. For example, in a T-maze paradigm, repulsion should only occur in the central stem of the maze, which is shared across the left-and right-turn routes. To our knowledge, rodent studies have not directly compared population-level neural similarity during overlapping vs. non-overlapping segments of a maze. Third, repulsion should be relatively slow to develop as it is inherently a learning phenomenon (Hulbert and Norman, 2014), which contrasts with the idea that coding properties of the hippocampus allow for an immediate orthogonalization of activity patterns. Finally, as an extension of the prediction that event overlap triggers divergence, we also conducted a novel analysis in which we tested whether the degree of learning-related plasticity that an individual hippocampal voxel experienced was predicted by initial representational overlap within that voxel. This allowed us to determine whether learning-related plasticity preferentially occurs in representational units that are shared across events (Norman et al., 2006;Norman et al., 2007).

## RESULTS

### Behavioral measures of route discrimination

In an initial behavioral experiment, subjects studied sets of real-world routes that traversed the New York University campus. For each subject, the set of routes included pairs that shared a common path before diverging to terminate at distinct destinations (‘overlapping routes’) and pairs with no paths in common (‘non-overlapping routes’) (Figure 1A). Importantly, each route contributed to both conditions. For example, ‘route 1’ and ‘route 2’ were overlapping routes, but ‘route 1’ and ‘route 3’ were non-overlapping routes (Figure 1C). Each route contained an initial segment that was shared with another route (Segment 1), and a later segment, including the destination, that was route-specific (Segment 2; Figure 1A). Although the real-world spatial locations of the overlapping segments were identical, the pictures for each route were taken at different times and therefore differed subtly in terms of pedestrians, vehicles, etc. (Figure 1C and Supplementary Videos 1-8). Routes were studied twice per round for 14 rounds. Subjects were instructed to learn each route (i.e., the specific path to each destination), but were not told the destination at the start of the route. After each study round, subjects were shown individual pictures drawn from the routes and were asked to select the destination associated with each picture. Of central interest was accuracy for pictures drawn from Segment 1 of each route because selecting the correct destination for these pictures required discriminating between overlapping routes. Overall, subjects selected the correct destination (‘target’) at a higher rate than the destination of the overlapping route (‘competitor’) (F_1,21_ = 43.31, p = 0.000002; Figure 2A). The relative rate of target vs. competitor responses also markedly increased across learning rounds (F_1,21_ = 38.11, p = 0.000004; Figures 2B and 2C).

**Figure 1.**
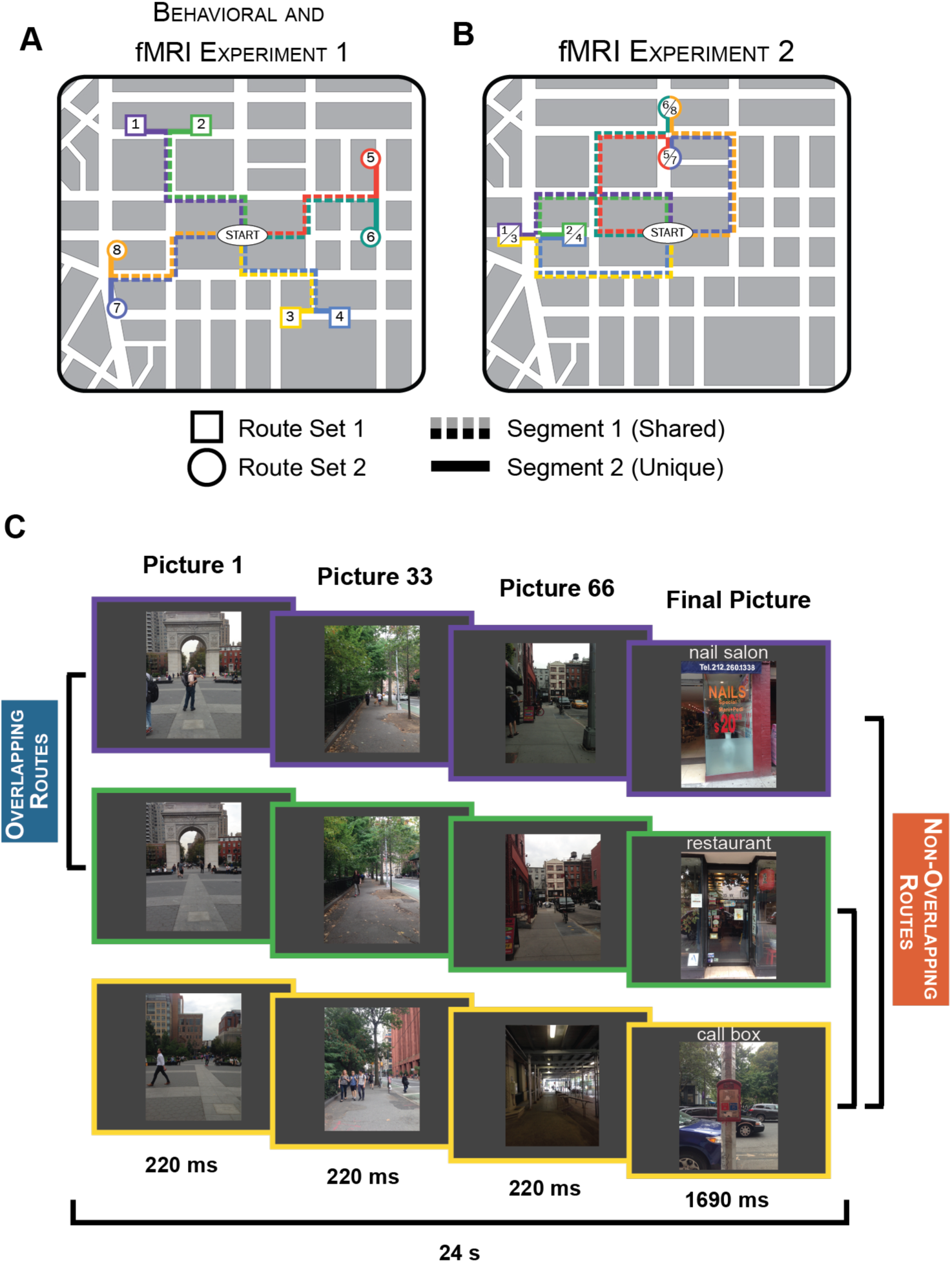
Route stimuli and experimental design. (A) In the behavioral experiment and fMRI Experiment 1, stimuli consisted of 8 routes that traversed the New York University campus. Each subject learned 4 routes–either Set 1 (routes 1-4) or Set 2 (routes 5-8). Each set included pairs of routes that shared a common path (‘overlapping routes’; e.g. routes 1 and 2) and pairs of routes with no common paths (‘non-overlapping routes’; e.g. routes 1 and 3). Individual routes contained two segments: Segment 1 refers to the portion of each route that shared a path with another route; Segment 2 refers to the unique portion of each route (after the overlapping routes diverged). (B) In fMRI Experiment 2 the stimuli again consisted of 8 routes with each subject learning 4 of the 8 routes, with the 4 routes per set containing overlapping and non-overlapping pairs. However, some of the non-overlapping route pairs in Experiment 2 terminated at the same destination (e.g. routes 1 and 3) whereas others terminated at distinct destinations (e.g., routes 1 and 4). (C) In each Experiment, each trial consisted of a series of rapidly presented pictures that lasted a total of 24s.

**Figure 2.**
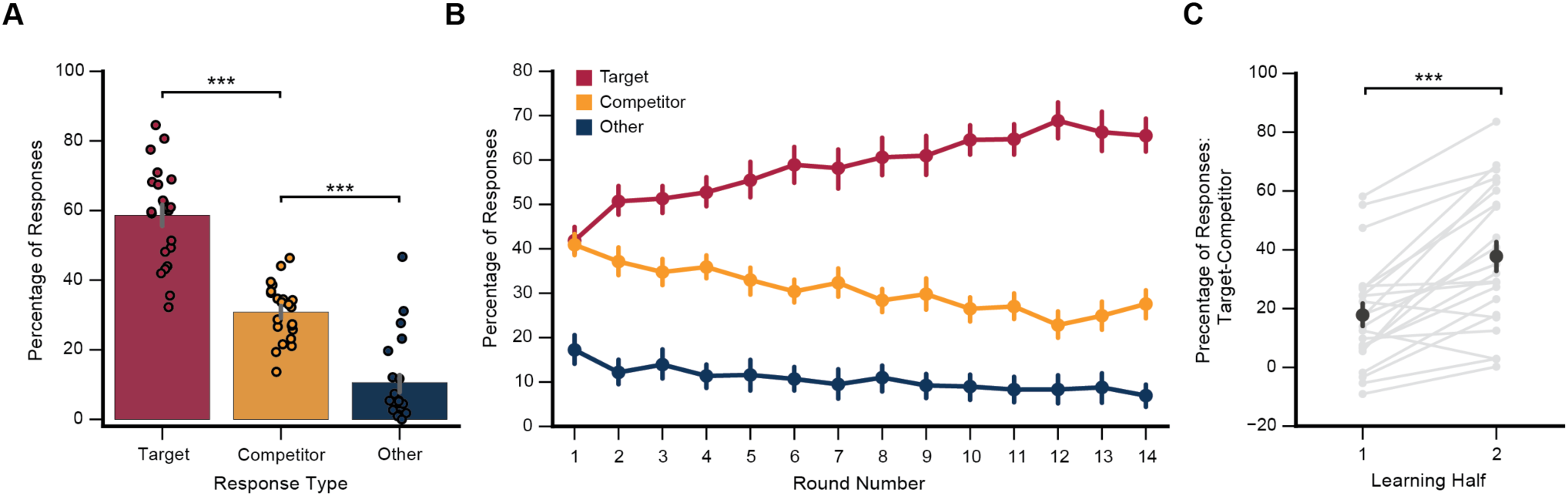
Memory performance for Segment 1 pictures in the behavioral experiment. (A) After each learning round subjects were shown static images sampled from each route and were asked to choose the corresponding destination from a set of four picture options: the target destination, the destination associated with the overlapping route (‘competitor’) and two destinations associated with non-overlapping routes (‘other’). Subjects were significantly more likely to select the target destination than competitor destination (F_1,21_ = 43.31, p = 0.000002) and significantly more likely to chose the competitor destination than other destinations (F_1,21_ = 41.39, p = 0.000002), despite the fact that ‘other’ options were more prevalent (2/4) than competitor options (1/4). (B) The relative percentage of target vs. competitor responses markedly increased over learning rounds (F_1,21_ = 38.11, p = 0.000004). (C) Discrimination between overlapping routes (percentage target responses - competitor responses) was significantly greater in the 2nd half of learning than the 1st half (t_21_ = 5.78, p = 0.00001). Error bars reflect 95% confidence intervals. *** p < 0.001

### Hippocampal representations of overlapping routes diverge with learning

We next tested for evidence of hippocampal repulsion of overlapping routes in two fMRI studies. The first fMRI study used the same stimuli as the behavioral study (Figure 1A). The second fMRI study used a new set of stimuli that again included overlapping and non-overlapping routes, but some of the non-overlapping routes terminated at a common destination (Figure 1B). Unless otherwise noted, all analyses below combine data across experiments and all comparisons of non-overlapping routes are restricted to those that terminated at distinct destinations. For Segment 1 of each route, we obtained a corresponding neural activity pattern by extracting voxel-wise patterns of activity as they unfolded over time. These spatiotemporal activity patterns were then correlated for every pair of routes, resulting in a correlation matrix reflecting pairwise route similarity (Figure 3A). We considered pattern similarity for (1) repetitions of the same route, (2) overlapping routes, and (3) non-overlapping routes. Separate correlation matrices were generated for each subject’s hippocampus and for a control region: the ‘parahippocampal place area’ (PPA), which is adjacent to the hippocampus and is involved in scene processing and navigation (Figure 3B) (Epstein et al., 1999;Epstein, 2008). Because our behavioral experiment indicated that discrimination of overlapping routes robustly improved from the 1st to 2nd half of learning (Figure 2C), we divided the fMRI data into halves and independently computed pattern similarity measures within each of these halves. As in the behavioral experiment, subjects in both fMRI experiments were able to successfully discriminate between the overlapping routes by the end of learning (see Figure S1).

**Figure 3.**
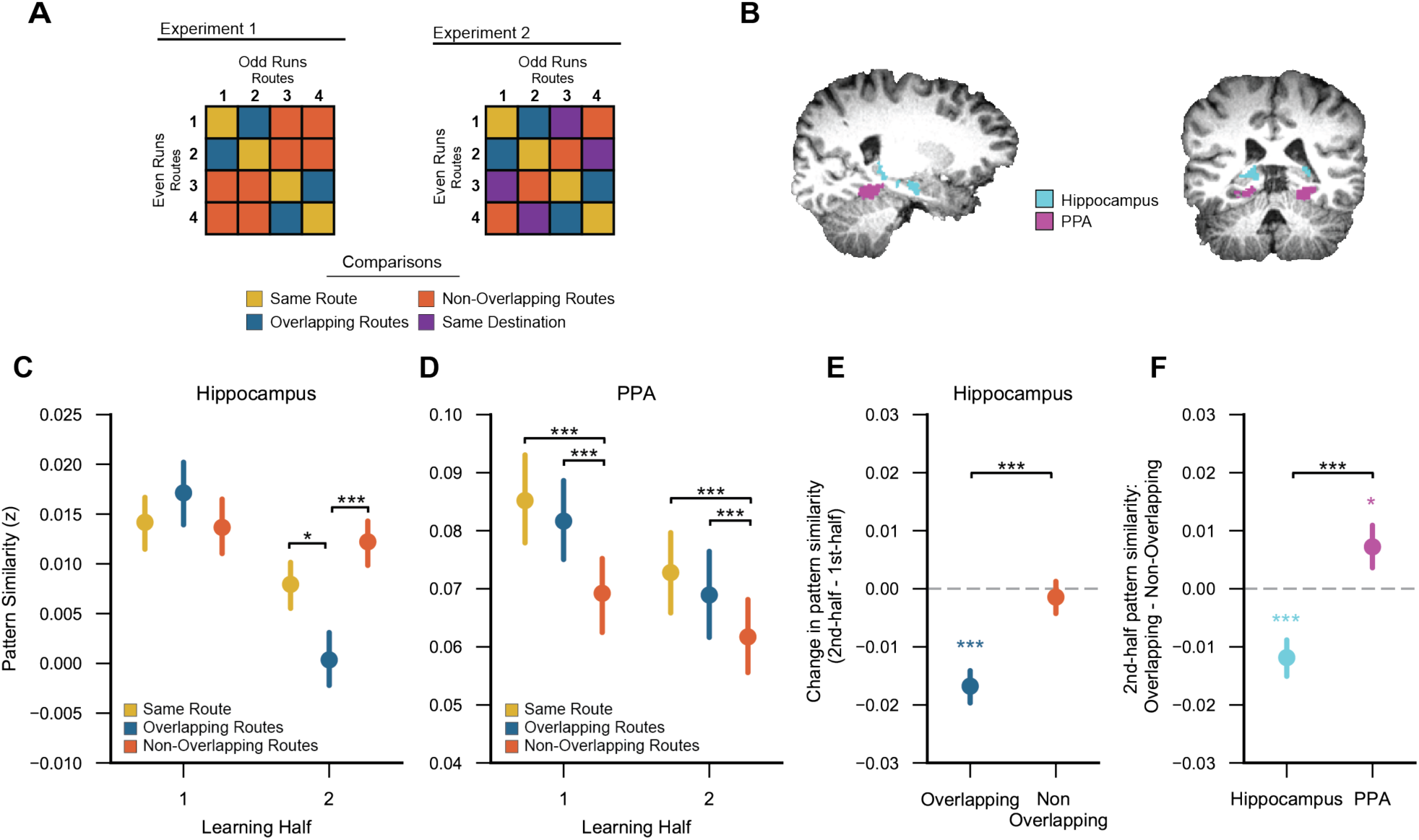
Learning-related changes in hippocampal pattern similarity. (A) Sample similarity matrices depicting analyses for Experiments 1 and 2 (routes 5-8 are not shown). For each Experiment, Pearson correlations were applied to spatiotemporal activity patterns to measure the similarity between: repetitions of the same route (‘same route’), routes with overlapping paths but distinct destinations (‘overlapping routes’), and routes with non-overlapping paths and distinct destinations (‘non-overlapping routes’). Experiment 2 included an additional comparison of routes with non-overlapping paths that ended at a common destination (‘same destination’). All correlations were applied to spatiotemporal activity patterns from independent fMRI runs (odd vs. even runs). (B) Hippocampus and parrahipocampal place area (PPA) regions of interest for a representative subject, displayed on T1 anatomical scan. (C) Within the hippocampus, the similarity of overlapping routes relative to same routes decreased across learning (1st vs. 2nd half; p = 0.009). Likewise, there was a learning-related decrease in the similarity of overlapping routes relative to non-overlapping routes (p = 0.0008). (D) Within PPA, there was no learning-related change in the relative similarity of overlapping vs. same routes (p = 0.96) nor in the relative similarity of overlapping vs. non-overlapping routes (p = 0.13). (E) Within the hippocampus, overlapping route similarity decreased across learning (1st vs. 2nd half, p = 0.0000006) whereas non-overlapping route similarity did not change with learning (p = 0.63). (F) In the 2nd half of learning, overlapping route similarity was significantly lower than nonoverlapping route similarity within the hippocampus (p = 0.0005; ‘reversal effect’) whereas in PPA the opposite was true: overlapping route similarity was significantly greater than non-overlapping route similarity (p = 0.038). Error bars reflect 95% confidence intervals. * p < 0.05, ** p < 0.01, *** p < 0.001.

Of critical interest, there was a learning-related decrease in pattern similarity among overlapping compared to non-overlapping routes, as reflected by an interaction between overlap (overlapping/non-overlapping) and learning (1st half/2nd half) (F_1,39_ = 13.163, p = 0.0008; Figure 3C). Whereas pattern similarity among overlapping routes decreased with learning (F_1,39_ = 35.21, p = 0.0000006), similarity among non-overlapping routes did not change (F_1,39_ = 0.24, p = 0.63; Figure 3E). This dissociation is striking when considering that all routes contributed to both the overlapping and non-overlapping comparisons. Thus, learning did not globally reduce similarity among routes; rather, learning specifically reduced similarity between overlapping routes. Moreover, overlapping route similarity decreased to the point that in the 2nd half of learning overlapping routes were markedly less similar than non-overlapping routes (F_1,39_ =14.20, p = 0.0005; Figure 3F). This result was significant in each of the fMRI experiments (ps < .05; see Figure S2 for results separated by experiment). Thus, despite the fact that overlapping routes were spatially and visually more similar than non-overlapping routes, the hippocampus represented overlapping routes as less similar than non-overlapping routes–a result we refer to as a ‘reversal effect’ because the representational structure is opposite to the inherent similarity structure of the routes. This reversal effect was not present in the 1st half of learning (F_1,39_ = 1.41, p = 0.24), confirming that it developed over learning (See Figure S3 for hippocampal pattern similarity computed at a finer time scale).

Next, we tested whether this reversal effect was selective to the overlapping route segments (Segment 1). If the reversal effect was triggered by route overlap, this makes a paradoxical prediction: that there should be relatively greater similarity among hippocampal activity patterns once the overlapping routes diverged. Indeed, when considering data from Segment 2–i.e., after overlapping routes diverged-the reversal effect was absent (F_1_,_39_ = 0.31, p = 0.58; Figures 4A and 4B). The selectivity of the reversal effect to the overlapping portion of the overlapping routes was confirmed by a significant overlap x segment interaction (2nd half data only, F_1,39_ = 4.28, p = 0.045). Thus, once overlapping routes diverged–and visual and spatial similarity decreased–hippocampal activity patterns became relatively more similar, indicating that the reversal effect was a reaction to overlap.

**Figure 4.**
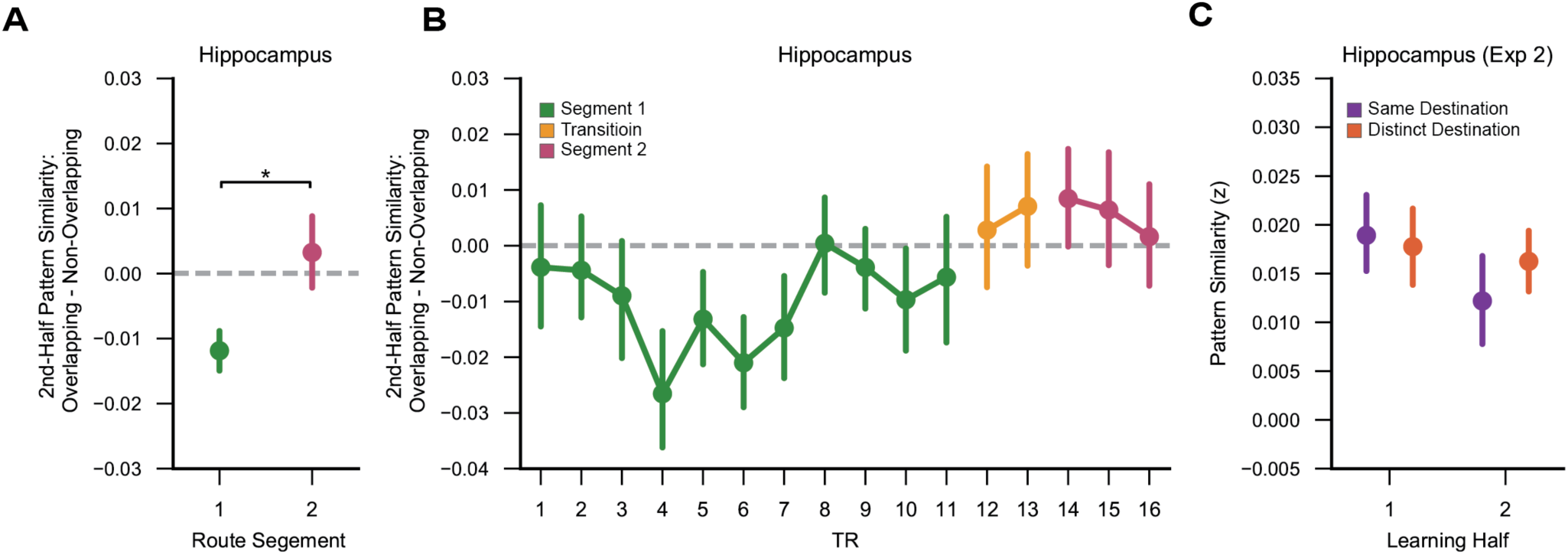
Hippocampal spatiotemporal pattern similarity across route segments. (A) In the 2nd half of learning, the hippocampal reversal effect (overlapping route similarity < non-overlapping route similarity) was significant for Segment 1 (p = 0.0005), but not Segment 2 (p = 0.58) and the interaction between overlap and segment was significant (p = 0.045). (B) TR-by-TR comparison of spatial pattern similarity for overlapping vs. non-overlapping routes. Spatial patterns analyzed at each TR were computed as the average pattern of a sliding 3-TR window. Transition TRs reflect time points that included the end of Segment 1 and the beginning of Segment 2. (C) In Experiment 2, non-overlapping routes included pairs of routes that terminated at distinct destinations as well as pairs that terminated at the same destination. Within Segment 1, there was no evidence for a learning-related change in spatiotemporal pattern similarity in the same destination condition relative to the distinct destination condition (p = 0.47). When considering the 2nd half data alone, there was also no difference in similarity between the same and distinct destination conditions (p = 0.34). Error bars reflect 95% confidence intervals. * p < 0.05.

Within PPA, there was no learning-related reduction in the similarity of overlapping compared to non-overlapping routes (Segment 1 data only; F_1,39_ = 2.42, p = 0.13; Figure 3D). In fact, overlapping route similarity was greater than non-overlapping route similarity in the 1st half (F_1,39_ = 21.01, p = 0.00005) and 2nd half of learning (F_1,39_ = 4.63, p = 0.038; Figure 3F; note: this effect differed across Experiments, see Figure S2). Thus, the inverted representational structure that was observed in the hippocampus by the end of learning was absent in PPA. The dissociation between the representational structure in PPA vs. hippocampus at the end of learning was reflected in a highly significant region x overlap interaction (F_1,39_ = 22.18, p = 0.00003). A similar dissociation was observed when comparing hippocampus to retrosplenial cortex, another region involved in scene-processing and navigation (see Figure S4).

Although we primarily focus on the comparison of overlapping vs. non-overlapping routes, the comparison of overlapping vs. same routes is also informative. To the extent that hippocampal representations of overlapping routes are distinct, overlapping route similarity should be lower than same route similarity (e.g, route 1 vs. route 2 similarity should be lower than route 1 vs. route 1 similarity). Indeed, within the hippocampus there was a learning-related change such that overlapping route similarity decreased relative to same route similarity (F_1,39_ = 7.59, p = 0.009). Overlapping route similarity was significantly lower than same route similarity in the 2nd half of learning (F_1,39_ = 5.61, p = 0.023), but not in the 1st half of learning (F_1,39_ = 0.85, p = 0.35). Within PPA, there was no learning-related change in overlapping vs. same route similarity (F_1,39_ = 0.003, p = 0.96). In PPA, the difference between overlapping and same route similarity was not significant in the 1st half (F_1,39_ = 0.89, p = 0.35) or 2nd half (F_1,39_ = 0.86, p = 0.36).

One way in which hippocampal route representations may diverge with learning is through the learned ability to predict the route destinations (Lee et al., 2006;Wikenheiser and Redish, 2015;Brown et al., 2016;Davachi and DuBrow, 2015; Ólafsdóttir et al., 2015;Hindy et al., 2016). To test this possibility we considered data from Experiment 2, which contained pairs of non-overlapping routes that terminated at distinct destinations as well as pairs of non-overlapping routes that terminated at the same destination. If hippocampal activity patterns reflected navigational goals, spatiotemporal pattern similarity from Segment 1 should be greater for non-overlapping routes that terminated at the same destination relative to non-overlapping routes that terminated at distinct destinations. However, we found no evidence for a difference between these conditions (Figure 4C). Specifically, there was no learning related increase in spatiotemporal similarity for same destination pairs relative to distinct destination pairs (F_1,20_ = 0.53, p= 0.47), nor was there a difference between same and distinct destination pairs when considering second-half data alone (t_20_ = 0.98, p = 0.34). Thus, the observed divergence of hippocampal activity patterns is not readily explained by destination coding.

### Voxel-Level changes in route similarity

The preceding results indicate that hippocampal representations of overlapping events–as reflected by distributed patterns of activity–diverged with learning, and that this divergence was triggered by route overlap. But what factors determined the level of plasticity that individual voxels exhibited? On the one hand, the reversal effect potentially reflects a global form of plasticity, with all voxels showing a comparable degree of learning-related divergence. However, a theoretically important alternative that is motivated by our main findings above is that the amount of initial representational overlap (within a voxel) determines the degree to which divergence occurs over the course of learning (Norman et al., 2006;Norman et al., 2007).

To test whether voxel-level divergence varied according to initial representational overlap, we characterized every voxel in terms of the similarity with which it responded to overlapping routes. However, because spatial pattern similarity cannot be computed at the level of individual voxels (i.e., a single voxel has no spatial pattern), we instead capitalized on the temporal dimension of our stimuli, computing the similarity of each voxel’s timecourse across pairs of routes. We refer to this measure as ‘timecourse similarity’ (Figure 5A). Voxels were rank-ordered by 1st-half timecourse similarity and binned into groups corresponding to ‘weak,’ ‘moderate,’ or ‘strong’ similarity (i.e., the bottom 1/3, middle 1/3, and top 1/3 of similarity values). Importantly, this binning was independently repeated for every pair of routes, each voxel in each region of interest, and each subject. Performing the analysis in a route-specific manner is important because a given voxel may exhibit strong timecourse similarity across one pair of routes but weak timecourse similarity across a different pair of routes. Timecourse similarity values from the 2nd half of learning were then obtained from these voxel bins. This allowed for timecourse similarity values at the end of learning to be expressed as a function of timecourse similarity at the beginning of learning. It is important to note that we did not measure changes in timecourse similarity from the first to second half, as such measures would be distorted by regression to the mean.

**Figure 5.**
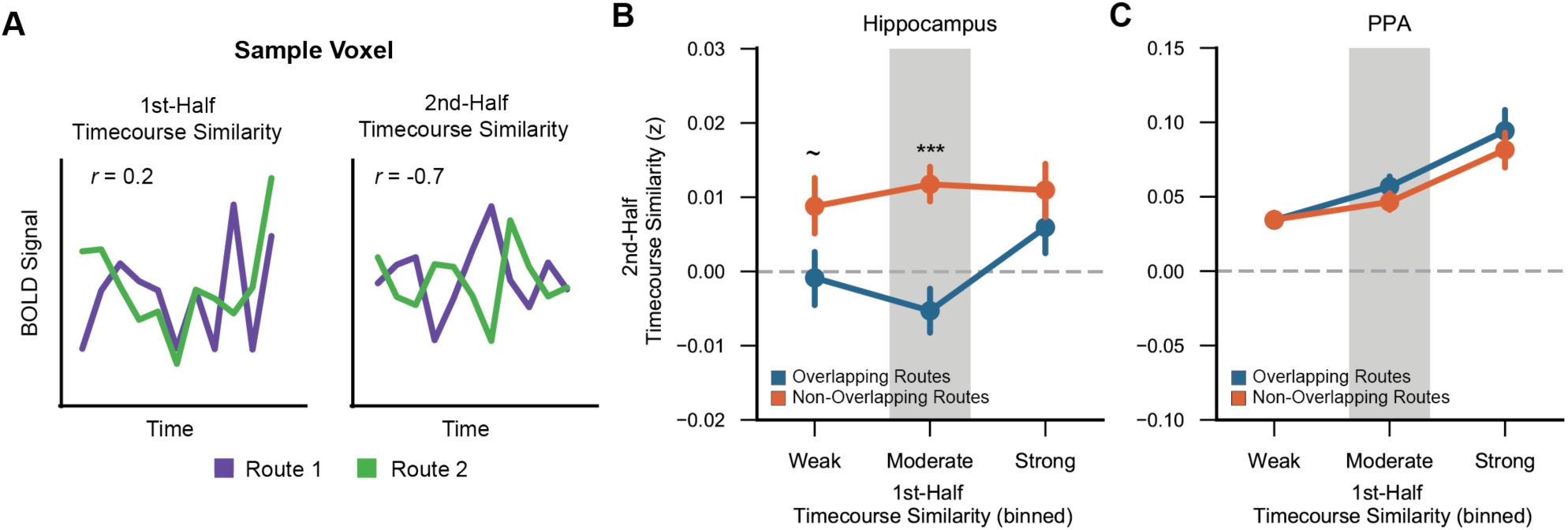
Voxel-level plasticity. (A) Timecourse similarity was defined as the correlation of a single voxel’s temporal pattern of activity across two different routes. For each voxel, timecourse similarity was separately computed for the 1st half and 2nd half of learning. (B,C) Second-half timecourse similarity plotted as a function of 1st-half timecourse similarity, separately for the hippocampus (B) and PPA (C) and for overlapping (blue) and non-overlapping (orange) routes. Within the hippocampus, 2nd-half timecourse similarity was markedly lower for overlapping than non-overlapping routes (reversal effect) for voxels that were moderately shared at the beginning of learning (p = 0.00009). ~ p < 0.1, *** p < 0.001.

Within the hippocampus, an ANOVA with factors of overlap (overlapping/non-overlapping) and bin (weak/moderate/strong) revealed a significant overlap x bin interaction (F_2,78_ = 3.19, p = 0.046). This interaction reflected a relatively greater difference between overlapping and non-overlapping routes (reversal effect) for voxels that exhibited ‘moderate’ timecourse similarity during the 1st-half of learning. Namely, the reversal effect was highly significant in the ‘moderate’ bin (F_1,39_ = 19.17, p = 0.00009), marginally significant in the ‘weak’ bin (F_1,39_ = 3.62, p = 0.064), and not significant in the ‘strong’ bin (F_1,39_ = 1.53, p = 0.22). Thus, the reversal effect was not consistent across voxels; rather, it was most pronounced among voxels that exhibited moderate similarity across overlapping routes at the beginning of learning. Considering overlapping routes alone–as opposed to the difference between overlapping and non-overlapping routes–2nd-half timecourse similarity also significantly varied according to 1st-half similarity (F_2,78_ = 4.74, p = 0.012), with the function qualitatively characterized by a dip for voxels in the ‘moderate’ bin (Figure 5B). Indeed, adding a quadratic term to a mixed-effects regression model that included a linear term significantly improved the model fit (*χ*^2^ = 6.06, p = 0.014), indicating a non-monotonic relationship between timecourse similarity at the beginning vs. end of learning. For non-overlapping routes, 2nd-half timecourse similarity did not vary according to 1st half similarity (F_2,78_ = 0.28, p = 0.76). See Figure S5 for the results of a complimentary Bayesian curve-fitting analysis that relates 1 st-half timecourse similarity to 2nd-half timecourse similarity.

The relationship between 1st and 2nd half timecourse similarity for overlapping routes was markedly different in PPA, as reflected by a significant region (hippocampus/PPA) x bin interaction (F_2,78_ = 18.12, p = 0.0000003). A region x bin x overlap interaction was marginally significant (F_2,78_ = 2.95, p = 0.0584). Qualitatively, PPA voxels that were moderately shared across overlapping routes in the 1st half of learning remained moderately shared in the 2nd half of learning (Figure 5C).

Collectively, these findings indicate that the hippocampal reversal effect preferentially occurred for voxels that were moderately shared at the beginning of learning. This suggests a ‘Goldilocks effect,’ wherein intermediate levels of overlap produced the strongest amount of learning-related divergence. At a more general level, this finding provides unique evidence that initial overlap among hippocampal representations is an important determinant of learning-related plasticity.

## DISCUSSION

Here, across two fMRI studies, we found that hippocampal representations of overlapping spatial routes dramatically diverged with learning–to the point that overlapping routes were coded as less similar than non-overlapping routes. This ‘reversal effect’ was clearly a result of learning as it was only evident after subjects gained considerable familiarity with the routes and it paralleled behavioral improvement in memory-based route discrimination. The result was also selective to the hippocampus, with no evidence of a reversal effect in PPA. Finally, using a novel analysis approach, we show that the learning-related divergence of hippocampal activity patterns was most pronounced for voxels that were moderately shared across overlapping routes at the beginning of learning.

### Measuring hippocampal disambiguation during human spatial learning

Although there is substantial evidence of hippocampal disambiguation of overlapping events in both human and non-human animals (Favila et al., 2016;Chadwick et al., 2010;Ginther et al., 2011;Agster et al., 2002;McKenzie et al., 2014;Frank et al., 2000;Kumaran and Maguire, 2006;Gilbert et al., 2001;Schlichting et al., 2015;Grieves et al., 2016;Kyle et al., 2015), there remains a gap between these two literatures as well as ambiguity concerning the underlying mechanisms. In rodent studies, hippocampal disambiguation has largely been measured in spatial navigation paradigms, with ‘T-maze’ paradigms being a common and elegant way to manipulate event overlap (Wood et al., 2000;Frank et al., 2000). In humans, however, most studies of hippocampal disambiguation involve static and usually arbitrary visual images with no spatial learning component (Favila et al., 2016;Kumaran and Maguire, 2006;Schlichting et al., 2015;Hsieh et al., 2014;Bakker et al., 2008;Hulbert and Norman, 2014). The present study represents an important step in bridging these literatures in that we adapted the canonical rodent T-maze paradigm to a rich and naturalistic human spatial learning paradigm. Thus, the present human fMRI study benefited from high ecological validity and much closer alignment with the paradigms that have traditionally been used to study hippocampal disambiguation in rodents. Because of the temporally dynamic nature of our route stimuli, we also measured hippocampal representations using a novel spatiotemporal pattern similarity analysis. This data-rich measure captures spatial patterns of activity (population-level codes) while preserving information about the temporal order in which these spatial patterns unfolded.

Several subtle details of our paradigm—and the analyses we employed—are critical for the interpretation of our results. First, our analysis approach specifically compared representations of overlapping events to representations of non-overlapping events (Favila et al., 2016). This allowed for learning-related changes to be expressed relative to a meaningful baseline—a baseline that, to our knowledge, is absent in rodent T-maze paradigms. Indeed, the fact that hippocampal representations of visually-and spatially-overlapping routes became less similar than routes that contained no spatial overlap or visual similarity is not only striking, but it provides essential insight into the underlying mechanism (a point we detail below). Second, our design did not involve separate sets of routes for the overlapping and non-overlapping comparisons (Kumaran and Maguire, 2006;Brown et al., 2010;Hsieh et al., 2014); rather, each route was included in each comparison. For example, whereas routes 1 and 2 represent overlapping routes, routes 1 and 3 represent non-overlapping routes. As such, any observed differences between overlapping and non-overlapping routes cannot be attributed to differences between the actual stimuli or to differences in attention, familiarity, vigilance, etc. It is also of note that our findings generalized across entirely different sets of stimuli (Experiments 1 and 2). Lastly, for our critical comparison of overlapping vs. non-overlapping routes, we focused on spatiotemporal activity patterns during the overlapping segments of the routes (Segment 1 data)–that is, before the overlapping routes diverged. Indeed, once the overlapping routes diverged (Segment 2 data), the hippocampal reversal effect ‘disappeared’ (Figures 4A and 4B). Thus, hippocampal representations of overlapping routes were most dissimilar when routes actually overlapped, clearly indicating that the reversal effect was triggered by event overlap.

### Mechanism underlying hippocampal reversal effect

While there is general agreement that the hippocampus disambiguates overlapping event representations—a phenomenon that has been termed ‘pattern separation’—there remains considerable debate about how pattern separation is achieved, with an emerging perspective that hippocampal pattern separation includes multiple computationally distinct mechanisms (Leutgeb et al., 2007;GoodSmith et al., 2017). However, the most prominent account is that pattern separation is achieved by sparse coding in the hippocampus—particularly within the dentate gyrus (O’Reilly and McClelland, 1994;O’Reilly and Rudy, 2001;Marr, 1971;Leutgeb et al., 2007;McHugh et al., 2007;Bakker et al., 2008;Yassa and Stark, 2011;Treves and Rolls, 1994;McClelland and Goddard, 1996;GoodSmith et al., 2017). With sparse codes, the probability of individual neurons being shared across representations is reduced and resulting representations are orthogonalized. While our data do not argue against the idea that sparse coding allows for orthogonalization, this account fails to explain our central findings. In particular, the maximum amount of separation that can be achieved by sparse coding is perfect orthogonalization, but the reversal effect we observed indicates that ‘over-orthogonalization’ occurred. That is, if every route is orthogonally coded, the similarity among overlapping routes will be precisely equal to—but not below—the similarity among non-overlapping routes. Thus, the present findings reveal a degree of pattern separation that goes beyond the typically assumed theoretical maximum.

An additional important consideration in understanding the observed reversal effect is that it emerged with learning. While there are several existing accounts of how learning contributes to the divergence of hippocampal activity patterns, most accounts again fail to explain the observed reversal effect. For example, the hippocampus is thought to play a critical role in establishing unique contexts for overlapping events (Hsieh et al., 2014;McKenzie et al., 2014). However, associating each route with a unique context should only reduce global similarity among events and does not explain why overlapping routes (which inherently share some contextual elements, such as location) would be less similar than non-overlapping routes. Similarly, hippocampal activity patterns may reflect predictions about route destinations (Lee et al., 2006;Wikenheiser and Redish, 2015;Brown et al., 2016;Davachi and DuBrow, 2015; Ólafsdóttir et al., 2015;Hindy et al., 2016), which could explain why hippocampal activity patterns diverge with learning, but cannot explain why hippocampal representations of overlapping events would diverge to the point of being less similar than non-overlapping events. Moreover, we did not observe any evidence of destination coding in the present study (Figure 4C), suggesting that, at least in the present study, hippocampal representations of the routes reflected information other than predicted destinations.

Conceptually, an appealing way to account for the hippocampal reversal effect is that route overlap triggered a repulsion of event representations (Norman et al., 2006; Norman et al., 2007;Hulbert and Norman, 2014). From this perspective, co-activation of similar memories triggered adaptive changes in hippocampal representations such that overlapping memories specifically ‘moved apart’ from one another. By analogy, this repulsion is similar to a teacher moving feuding children to opposite corners of a classroom in that the goal is to specifically increase the distance between the feuding children (as opposed to the distance between all children). Thus, in contrast to orthogonalization, where overlapping memories are represented as ‘unique,’ a repulsion account holds that overlapping memories are represented as ‘different from one another.’ A repulsion account is not only consistent with the observed reversal effect but also readily explains the striking and seemingly paradoxical finding that the hippocampal reversal effect ‘disappeared’ precisely once routes diverged (Segment 2; Figures 4A and 4B). The idea of repulsion among hippocampal representations has been elegantly described in biologically plausible computational models of the hippocampus, and the mechanism underlying this repulsion has been termed ‘differentiation’ (Hulbert and Norman, 2014;Kim et al, 2017). While a limited number of human fMRI studies have provided strong hints of differentiation in the hippocampus (Hulbert and Norman, 2014;Favila et al., 2016;Schlichting et al., 2015;Schapiro et al., 2012;Kim et al, 2017), the present findings provide the strongest and most unambiguous evidence to date that hippocampal representations of overlapping events diverge to the point that they are less similar than non-overlapping events.

Because we measured hippocampal similarly over the course of an extended learning paradigm, we were also able to provide insight into the timecourse of these representational changes. Importantly, we show that the reversal effect was remarkably slow to emerge—with evidence of the reversal effect only emerging after routes had been presented approximately 20 times (see Figure S3). However, this slow timecourse strongly paralleled the timecourse of behavioral improvements in memory-based discrimination of the overlapping routes, as identified in a separate behavioral study (Figure 2). The parallel between the timecourse of behavioral improvement and the hippocampal reversal effect is consistent with the idea that differentiation is a learning-related phenomenon (Hulbert and Norman, 2014;Kim et al, 2017) and that differentiation is behaviorally relevant (Favila et al., 2016).

One interesting and potentially surprising aspect of our findings is that during the initial stages of learning, hippocampal representations of overlapping routes were numerically but not significantly more similar than non-overlapping routes (Figure 3A). On the one hand, this may reflect insensitivity of our measurements to a small but real difference. On the other hand, the lack of initial difference between overlapping and non-overlapping routes may reflect an immediate orthogonalization of route representations in the hippocampus (O’Reilly and McClelland, 1994;O’Reilly and Rudy, 2001;Marr, 1971;Leutgeb et al., 2007;McHugh et al., 2007;Bakker et al., 2008;Yassa and Stark, 2011;Treves and Rolls, 1994;McClelland and Goddard, 1996;GoodSmith et al., 2017;Treves and Rolls, 1994). From this perspective, the slow-acting reversal effect we observed may have been driven by any residual representational overlap that remained after initial orthogonalization.

### Voxel-level plasticity

Although several fMRI studies—including the present study—have shown that hippocampal activity patterns change with learning, a separate and under-explored question is whether there are specific factors that predict the degree to which individual voxels will change with learning. Here, motivated by our primary findings that route overlap triggered repulsion of hippocampal representations, we considered whether the degree of overlap in individual voxel representations predicted the reversal effect (a measure of plasticity) that a voxel experienced. In most fMRI studies, this question would be difficult to address, because representational overlap, as indexed by spatial pattern similarity, cannot be computed at the level of a single voxel. Here, however, because of the temporally-dynamic nature of our stimuli, we were able to use timecourse similarity to measure the similarity with which a single voxel responded to each pair of routes. In this manner, we quantified the strength of the reversal effect within individual voxels. Indeed, we observed that the reversal effect was not evenly distributed across voxels; rather, there was a ‘sweet spot,’ with the reversal effect disproportionately occurring in voxels that exhibited ‘moderate’ degrees of timecourse similarity at the beginning of learning.

Why might the reversal effect disproportionately occur for voxels with moderate levels of initial timecourse similarity? To answer this, it is first important to consider what timecourse similarity reflects. When a voxel responds similarly to a pair of overlapping routes (i.e., high timecourse similarity), this suggests that the voxel–or ensembles of neurons within that voxel–are ‘shared’ across those routes’ representations. Critically, it is proposed that this form of representational ‘sharing’ is precisely what triggers hippocampal differentiation. Namely, if two overlapping events—A and A’—share common representational units (voxels, neurons, or connections between neurons), then activation of one event (A) is likely to activate the overlapping event (A’), and vice versa. For example, when viewing route 1, it is likely that route 2 (the overlapping route) is partially activated, owing to shared representational units (Tanaka et al., 2014;Cai et al., 2016;Kuhl et al., 2011). When this occurs, the co-activated representation is subject to plasticity. Interestingly, and central to interpreting the present findings, it is argued that the plasticity that these co-activated units experience is non-monotonically related to their level of activation, with moderately activated units subject to weakening, whereas strongly activated units are strengthened and weakly activated units do not experience plasticity (Norman et al., 2006;Norman et al., 2007;Newman and Norman, 2010;Detre et al., 2013;Poppenk and Norman, 2014;Lewis-Peacock and Norman, 2014;Hulbert and Norman, 2014). Putatively, this non-monotonic plasticity rule reflects a competition between excitation and inhibition, with moderate activation corresponding to inhibition ‘overcoming’ excitation. From this perspective, the present finding of a non-monotonic relationship between initial timecourse similarity and the reversal effect potentially reflects the same putative non-monotonic relationship between activation and plasticity. That said, our analysis does not constitute a direct test of this model—mainly because timecourse similarity is not a direct measurement of co-activation. However, this perspective offers a theoretically grounded and biologically plausible interpretation of our findings. Regardless of the specific mechanistic account, the present findings provide novel evidence that the degree of representational divergence experienced by individual hippocampal voxels is determined, at least in part, by the degree of representational overlap during initial stages of learning. This finding further strengthens our argument that overlap itself triggers a repulsion of hippocampal representations.

### Nature of hippocampal representations

A final question concerns the nature of information coded for by the hippocampus. While the hippocampus is part of a broader network of regions involved in spatial navigation and memory (Spiers and Barry, 2015;Burgess et al., 2002;Ekstrom et al., 2003;Doeller et al., 2008), regions within this network may code for qualitatively different information. Indeed, we found that, by the end of learning, PPA–a cortical region adjacent to the hippocampus that has been implicated in coding spatial landmarks (Marchette et al., 2015)–exhibited a representational structure that was opposite to the representational structure in the hippocampus. Namely, in PPA overlapping routes were more similar than non-overlapping routes, whereas in the hippocampus this representational structure was reversed. As far as specific kinds of information that the hippocampus may have encoded, our findings do not, as noted above, suggest that the hippocampus prospectively coded for route destinations (Grieves et al., 2016). Moreover, it is not clear whether hippocampal activity patterns in the current study reflected current spatial locations (Miller et al., 2013;Doeller et al., 2008;Hassabis et al., 2009;Nielson et al., 2015;Ekstrom et al., 2003). Indeed, at first pass our findings appear to be at odds with a spatial coding account in that hippocampal activity patterns specifically diverged when routes were spatially overlapping. However, an intriguing, though speculative, possibility is that route overlap elicited the formation of separate spatial reference frames for each route (Kentros et al., 1998;Leutgeb et al., 2005;Bostock et al., 1991). In other words, hippocampal activity patterns in the present study may have reflected spatial codes so long as overlapping routes were represented within distinct maps. Alternatively, the hippocampus may have coded for non-spatial information that differentiated the overlapping routes (e.g., pedestrians, cars, etc.). Although these questions are beyond the scope of the present study, a clear priority in future studies is to better characterize how the divergence of hippocampal activity patterns corresponds to changes in the information encoded in these patterns.

## Author Contributions

A.J.H.C, S.E.F. and B.A.K designed the experiment. A.J.H.C. and A.O. ran the experiment. A.J.H.C. analyzed the data. A.J.H.C., S.E.F., and B.A.K. wrote the paper.

## Acknowledgements

We thank Anthony Stigliani and Kalanit Grill-Spector for providing stimuli for the category localizer. This work was supported by a grant from the National Institutes of Health (1RO1NS089729) to B.A.K.

## METHODS

### Subjects

New York University (NYU) students and alumni who were familiar with the NYU campus participated in the study. Subjects were restricted to NYU alumni and students in order to facilitate route learning and to reduce potential between-subject variance. Subjects were between the ages of 18-35, right-handed, native English speakers, had normal or corrected-to-normal vision and had no history of neurological disorders. Twenty-two subjects participated in the behavioral experiment (15 female; mean age = 20.77). Two additional subjects’ data were not collected due to technical errors. Twenty subjects (13 female; mean age = 22.15) participated in fMRI Experiment 1. Four additional subjects were excluded from data analysis - one for falling asleep in the scanner, two for technical errors during scanning, and one due to unreliable localizer data (see Regions of Interest). Twenty-one subjects (9 female; mean age = 23.17) participated in fMRI Experiment 2. One additional subject’s data was excluded from data analysis due to excessive head motion and another additional subject was excluded for technical errors during scanning. Sample sizes for the fMRI studies were based on a similar experiment from our lab (Favila et al., 2016) Informed consent was obtained according to procedures approved by the New York University Committee on Activities Involving Human Subjects.

### Stimuli and Design

In the behavioral experiment and fMRI Experiment 1 the stimuli consisted of eight routes that traversed the NYU campus (Figure 1A). Each route was comprised of a series of 98 unique pictures. All pictures were taken at regular intervals (every 10 paces) from an egocentric perspective by a researcher walking along the route. All routes started in the same location and made exactly three turns before ending at distinct destinations. Critically, the 8 routes consisted of 4 overlapping pairs. Overlapping pairs followed the same path for the majority of the route before diverging on the third turn to their respective destinations. The pictures for each route were taken at different times and therefore the pictures during the overlapping portion of routes were subtly different and could be distinguished from one another based on subtle differences in the pedestrians, vehicles, lighting, etc. For analysis purposes, routes were divided into pairs that shared an overlapping path (‘overlapping routes’; e.g. routes 1 and 2) or took distinct paths (‘non-overlapping routes’; e.g. routes 1 and 3). Furthermore, each route was divided into two segments: ‘Segment 1’ refers to the segment of each route that overlapped with another route and ‘Segment 2’ refers to the route-unique segment of each route. The third turn–which marked the boundary between Segments 1 and 2– occurred at the exact same picture numbers within pairs of overlapping routes (e.g., for routes 1 and 2) and varied minimally (between picture numbers 74-77) across sets of overlapping pairs (e.g., for routes 1/2 vs. routes 3/4). Likewise, all turns within a pair of overlapping routes occurred at identical time points in order to maximize the similarity of overlapping routes. There was exactly one overlapping pair that left the starting point in each cardinal direction (north, south, east, west). The 8 routes were divided into 2 sets (north/south routes and east/west routes). Each subject was assigned one set of routes (4 routes total) to learn, with the assignment of route sets alternating subject-by-subject. We included 2 sets of routes in order to ensure our results could not be explained by the idiosyncrasies of any one route.

A new set of 8 routes was used in fMRI Experiment 2 (Figure 1B). The routes were constructed using the same parameters as the routes used in the behavioral and first fMRI experiments, with one key difference. Instead of all routes terminating at distinct locations, fMRI Experiment 2 contained pairs of routes that took distinct paths but ended at the same destination. As before, the 8 routes were divided into two sets of 4 and each set of 4 contained two pairs of overlapping routes. The routes in each set could be divided into pairs that (a) shared an overlapping path but terminated at distinct destinations (‘overlapping routes’; e.g. routes 1 and 2), (b) had non-overlapping paths and terminated at distinct destinations (‘non-overlapping routes’; e.g. routes 1 and 4) or (c) had non-overlapping paths but terminated at the same destinations (‘same destination’; e.g. routes 1 and 3). Due to geographical constraints, the third turn (i.e. when overlapping routes diverged) in this set of routes occurred slightly later (between picture numbers 84-86) than in the set used in the behavioral and first fMRI experiments.

Videos of the overlapping route pairs used in the experiments are available in Supplementary Videos 1-8.

### Procedure

#### Behavioral Experiment

##### Route Learning

Subjects completed 14 rounds of route learning, with each route presented twice per round in random order. During a route learning trial, pictures from a route were presented in rapid succession (220 ms per picture, 10 ms blank screen in between pictures). Importantly, subjects were not told the destination of the route prior to the trial. Rather, the destination was only revealed at the end of the route, with the final picture (the destination) presented for 1690 ms. The destination’s name was also displayed above the final picture. Each route learning trial lasted a total of 24s and was followed by a 1-s inter-trial interval (ITI) during which a fixation cross was presented. Each round also contained two ‘catch’ trials to ensure subjects’ vigilance but were excluded from all analyses. For each catch trial, a route began as with a normal trial but the presentation stopped at a pre-selected picture number. A cue then appeared above the picture either instructing participants to identify (1) the routes’ final destination (destination test) or (2) the direction of the next turn (direction test). During the 3s response period the picture and test cue remained on screen with the four destination labels (destination test) or left/right labels (direction test) printed below the picture and participants selected their response using a keyboard. Catch trials stopped on pictures presented between 3-15s after the trial onset and at intervals of 1.5 s (to coincide with the TR length in the fMRI experiments; see fMRI Acquisition). The combined duration of the two test trials within each round were constrained to equal the duration of a full route learning trial (24 seconds). Although each subject completed an equal number of destination and direction catch trials throughout the experiment, and each route was tested an equal number of times, the assignment of catch trial type to both route number and round was randomized so as not to be predictable. That is, within a given round there could be 2 destination catch trials, 2 direction catch trials, or 1 of each, and a given route could be tested twice via a destination catch trial, twice as a direction catch trial, or once as each test.

##### Inter-Round Picture Test

At the end of each of the 14 route learning rounds, subjects were shown 20 static pictures, one at a time, drawn from the routes (5 per route in random order) and for each picture subjects were asked to select the corresponding destination from a set of four label options. The inter-round picture test was self-paced and subjects responded via keyboard. To ensure that the five pictures tested from each route in each test round were evenly sampled across positions in the route, each route’s 95 pictures (excluding the last 3 pictures that contained visuals of the destination) were divided into 5 time-bins of 19 pictures. For each inter-round picture test, one picture from each time-bin, from each route, was randomly selected to be tested with the constraint that a given picture was only tested once throughout the experiment. Responses on the test were divided into three groups: (1) ‘target’ if subjects selected the correct destination, (2) ‘competitor’ if subjects selected the overlapping route’s destination, and (3) ‘other’ if subjects selected the destination from a non-overlapping route.

##### Map Test

In order to assess each subject’s knowledge of the routes, subjects also completed a map test after finishing all rounds of route learning. For each trial on the map test, subjects were cued with a picture of a route’s destination for 4s. A map of the NYU campus then appeared on screen and subjects had 8s to click on the spatial location of the cued destination using a computer mouse. They were then prompted to draw with a pen the route taken to that destination on a paper print out of the campus map. Finally, participants completed both the Santa Barbara Sense of Direction Scale (SBSOD) and the Questionnaire on Spatial Representation (QSR) to assess their spatial acuity and reasoning. Results from the map test and questionnaires are not reported in the current study.

#### fMRI Experiments 1 and 2

##### Route Learning

The procedures from the behavioral experiment were slightly modified to be suitable for fMRI scanning. In both fMRI experiments, subjects first completed 2 practice route learning rounds (2 repetitions of each route per round) to familiarize them with the routes and task structure. Subjects then entered the scanner and completed an additional 14 rounds of route learning. The practice rounds were identical to the scanner rounds except that the first practice round did not contain any catch trials. During the scanned route learning rounds, the ITI was 6s (fixation cross) to allow for better separation of the hemodynamic response.

##### Inter-Round Picture Test

The inter-round picture test used in the fMRI experiments was shorter than in the behavioral experiment. In the fMRI version, there were a total of only 4 trials which contained pictures randomly sampled from the 4 routes. The sampled pictures were not constrained to be from different routes. The only constraint was that the pictures used in the inter-round picture test were not used in the post-scan memory test (described below). Additionally, in the fMRI version of the inter-round picture test subjects were shown each picture for a fixed amount of time (2.5 s) and could only respond during that time, using an MRI-compatible button box. Because the inter-round picture tests in the fMRI experiments only sparsely assessed route learning, these data are not reported. These test trials were only included to motivate subjects to learn the routes.

##### Functional Localizer

Following the 14 rounds of route learning subjects completed one localizer scan that was used to functionally define regions of interest for the fMRI analyses. The localizer scan contained 36 alternating blocks of three image types (12 blocks per category): faces, scenes (hallways or houses), and objects (cars or guitars). Each block lasted a total of 6s and contained 12 greyscale images presented for 500ms each. Subjects pressed a button whenever they detected a scrambled image, which occurred on half of all blocks (counterbalanced across category). An additional 12 baseline ‘blocks’ showing a blank grey screen (also 6s each) were randomly interspersed with the other blocks.

##### Post Tests

After exiting the scanner subjects first completed a map test (identical to the behavioral experiment). Next, subjects completed an extended picture test which included ten pictures drawn from each route (every 10th picture from picture 4 to 94), tested in random order. On each trial, the route picture was presented above the set of destination names (4 destination names in Experiment 1 and 2 destination names in Experiment 2). Subjects used a computer mouse to click on the destination name associated with each picture. This test was self-paced. Finally, subjects completed the Santa Barbara Sense of Direction Scale (SBSOD) and the Questionnaire on Spatial Representation (QSR).

### fMRI Data Analysis

#### MRI Acquisition

Scanning was performed on a 3T Siemens Allegra head-only scanner at the Center for Brain Imaging at New York University using a Siemens head coil. Structural images were collected using a T1-weighted protocol (256 × 256 matrix, 176 1-mm sagittal slices). Functional images were acquired using a T2^*^ weighted EPI single shot sequence containing 26 contiguous axial slices oriented parallel to the long-axis of the hippocampus (repetition time = 1.5 s, echo time = 23 ms, flip angle = 77 degrees, voxel size = 2 × 2 × 2 mm). The functional images did not cover the entire brain; rather, a limited field of view centered on the hippocampus was chosen in order to improve spatial resolution of data from the hippocampus. For the route learning scans, the first 6 volumes (during which time a "Get Ready" screen was presented, followed by a fixation cross) were discarded to account for T1 stabilization. For the localizer scan, the first 8 volumes and last 8 volumes (during which time a fixation cross was presented) were discarded. Field map and calibration scans were collected to improve functional-to-anatomical coregistration.

#### fMRI Preprocessing

Images were preprocessed using SPM8 (Wellcome Department of Cognitive Neurology, London, United Kingdom), FSL (FMRIB’s Software Library, Oxford, United Kingdom) and custom Matlab (The MathWorks, Natick, MA) routines. The preprocessing procedures included correction for head motion, coregistration of functional to anatomical images (using a registration procedure that aligned both functional and anatomical images to a calibration scan), and an unwarping procedure. Images from the functional localizer scan were spatially smoothed using a 4-mm full-width/half-maximum Gaussian kernel. Images from the route learning phase, which were used for pattern analyses, were smoothed using a moderate 2-mm full-width/half-maximum Gaussian kernel in order to improve signal-to-noise ratio. Prior research suggests that smoothing does not reduce sensitivity of pattern-based fMRI analyses (Op de Beeck, 2010). All analyses were performed in subjects’ native space.

#### fMRI univariate analysis

To analyze the localizer data, SPM was used to construct a general linear model with three regressors of interest corresponding to the three visual categories (scenes, faces, objects). These regressors were constructed as boxcar functions that onset at the first image of a category block and lasted for the duration of the block. Motion, block, and linear drift were modeled as regressors of no interest. All regressors were convolved with a canonical double-gamma hemodynamic response function. A linear contrast of scenes vs. faces and objects was used to obtain voxelwise estimates of scene sensitivity and a linear contrast of faces, scenes, and objects vs. baseline was used to obtain voxelwise estimates of visual sensitivity.

#### Regions of interest

Anlyses were performed using a region of interest (ROI) approach targeting the hippocampus, parahippocampal place area (PPA), and retrosplenial cortex (RSC). Anatomical hippocompal ROIs were defined using freesurfer’s automated subcortical segmentation procedure. The resultant hippocampal ROIs were then visually inspected and manually edited for any inaccuracies before registering them to each subject’s functional space. In order to identify voxels with high signal-to-noise ratios and to create ROI masks the same size as the PPA and RSC masks (see below), the hippocampal ROI consisted of the top 300 visually-responsive voxels within bilateral hippocampus, as determined from the category localizer (contrast of faces, scenes, and objects vs. baseline). Although this voxel selection procedure was implemented to increase our sensitivity to detect small differences in hippocampal patterns, it is important to note that our main findings were not dependent on such selection methods. Indeed when no voxel selection was applied within the hippocampus the interaction between overlap (overlap/non-overlap) and learning (1st half/2nd half) remained significant (F_1,39_ = 4.75, p = 0.0354), as did the reversal effect in the 2nd half of learning (F_1,39_ = 7.30, p = 0.0102).

PPA and RSC were identified using a combination of the category localizer and group-based probabilistic scene-selective ROIs identified from previous studies (Julien et al., 2012); http://web.mit.edu/bcs/nklab/GSS.shtml). First, the group-based probabilistic PPA and RSC masks were registered to each subject’s native space and voxels overlapping with the anatomically defined hippocampal masks were removed from the PPA/RSC masks to ensure independent ROIs. Then, the top 300 scene-selective voxels (contrast of scenes vs. faces and objects from the category localizer) within PPA and, separately, within RSC, were selected. This method ensured that the PPA and RSC ROIs were subject-specific but equal in size (number of voxels) and general location across all subjects (Marchette et al., 2015). Note: we chose 300 voxels as an a priori threshold for all our ROIs. This number corresponded to roughly the top 20% of the hippocampal voxels, 30% of the voxels within the group-based PPA mask, and 15% of the voxels in the group-based RSC mask. One subject from Experiment 1 was excluded because the average *t* value within their PPA ROI was more than two standard deviations below the mean PPA response in Experiment 1 (this was the only subject with a mean PPA or RSC response that was more than 2 standard deviations below the experiment mean); subjective assessment of data from this subject confirmed that there was no well-defined cluster within the group-based PPA mask that selectively responded to scenes.

#### Spatiotemporal pattern similarity

Pattern similarity analyses were performed on ‘raw’ (unmodeled) fMRI data. Several additional preprocessing steps were performed prior to performing pattern analyses. Functional images were detrended, high-pass filtered (0.01 Hz), and then z-scored within run. For route learning trials, volumes 3-19 (corresponding to 3-27s after stimulus onset) were divided into volumes corresponding to Segment 1 (i.e. the portion of each route that shared a path with another route) and Segment 2 (i.e. the unique portion of each route after overlapping paths diverged). The volume in each route corresponding to the transition between Segments 1 and 2 (i.e., the third turn in the routes) was discarded from analyses in order to keep Segments 1 and 2 distinct. In Experiment 1, Segment 1 occurred within the first 11 volumes and Segment 2 occurred within the last 4 volumes. In Experiment 2 the overlapping routes diverged slightly later; thus, Segment 1 corresponded to the first 12 volumes Segment 2 corresponded to the last 3 volumes. To perform pattern analyses, spatial activity patterns were concatenated across volumes of interest so that each route Segment was represented by a spatiotemporal pattern of activity whose vector length was equal to the number of voxels within an ROI x the number of TRs included in the Segment.

For each subject and each ROI, we computed pattern similarity scores (Pearson correlations) reflecting the representational similarity across each pair of routes. Correlations were always performed using data from independent fMRI runs (odd and even runs) in order to ensure independence. Thus, for analysis of data from the first half of learning, each route’s average spatiotemporal activity pattern was obtained from runs 1, 3, and 5 (odd runs) and, separately, from runs 2, 4, and 6 (even runs); average ‘odd run patterns’ were then correlated with average ‘even run patterns.’ Likewise, for analysis of data from the second half of learning, each route’s average spatiotemporal activity pattern was obtained from runs 9, 11, and 13 (odd runs) and, separately, from runs 10, 12, and 14 (even runs), and odd and even patterns were correlated. Data from runs 7 and 8 were excluded in order to ensure an equal number of odd and even runs within each half. Because each subject studied 4 routes, a 4 × 4 correlation matrix was generated for each subject (Figure 3A). Before any correlation values were averaged within conditions (e.g., overlapping routes), correlation coefficients were z-transformed (Fisher’s z).

#### Timecourse similarity

Timecourse similarity indexed the degree to which individual voxels were ‘shared’ across a given pair of routes. To compute timecourse similarity, we first obtained route-specific vectors of activation (using Segment 1 data only) for each voxel. The length of each timecourse vector was equal to the number of Segment 1 TRs (11 in Experiment 1; 12 in Experiment 2). Timecourse vectors were separately averaged across odd and even runs within each half (as with the spatiotemporal pattern analyses). Average timecourse vectors were then correlated (Pearson correlation) for every pair of routes, separately for each learning half (Figure 4B). Resulting correlation coefficients were z-transformed (Fisher’s z).

### Statistics

For all behavioral and fMRI analyses we used standard random-effects statistics (paired sample *t*-tests and repeated measures ANOVA). Two-tailed tests were used throughout at an alpha threshold of 0.05. Unless otherwise noted, analyses combined data across Experiments 1 and 2. For all ANOVAs run on these combined data, experiment number was included as a factor. For all of the hippocampal ANOVA effects described in the main text, interactions with experiment number were not significant (*Ps* > 0.2). See Figure S2 for hippocampal and PPA data separated by experiment. Mixed-effects regression models were used to assess the shape of the function relating timecourse similarity measures across experimental halves and were implemented in the lme4 package for R (http://lme4.r-forge.r-project.org). All models were constructed with random intercepts for each subject.

### Data and Code Availability

Raw data from the experiment is available on OpenFMRI (https://openfmri.org/dataset/ds000217) and code to run the analyses are available upon request from the first author.

**Figure S1.**
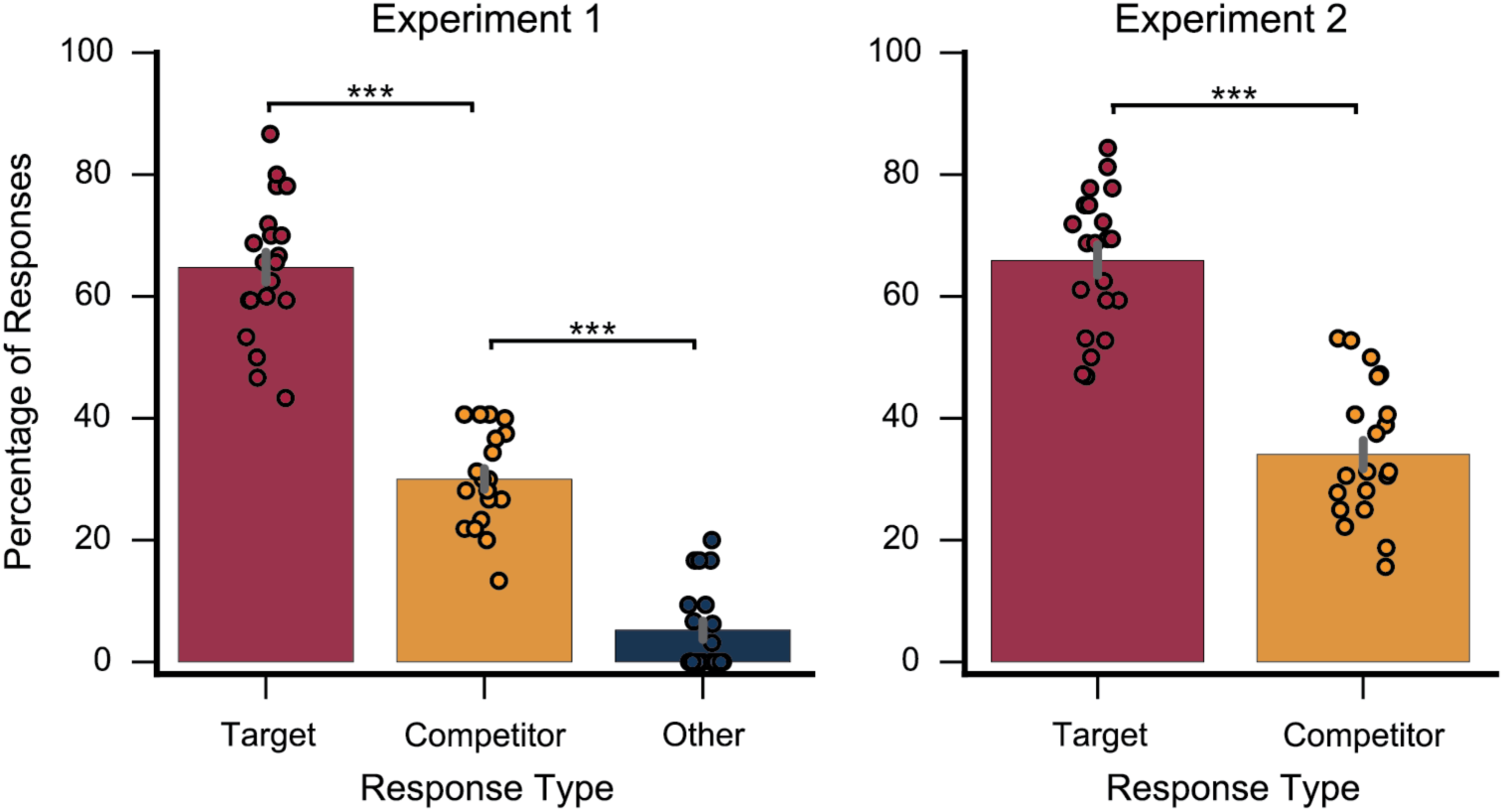
Related to *Figure 2*. Behavioral results from the post-scan picture test in each fMRI Experiment. After finishing all 14 learning rounds and the localizer scan subjects exited the scanner and completed the picture test. On each trial, subjects were shown a static picture drawn from one of the route stimuli (see Methods). Directly below each picture was a set of destination names. Subjects were instructed to select the destination name corresponding to the route picture. In Experiment 1, each trial had 4 destination options corresponding to: the target destination; the overlapping route destination (‘competitor’); and the two non-overlapping route destinations (‘other’). In Experiment 2 all of the routes studied by each subject ended in one of two possible destinations. Therefore, on each trial, the two destination options corresponded to either the target destination or the overlapping route destination (‘competitor’). Analyses were restricted to pictures drawn from Segment 1 of each route (i.e., the segments that contained overlap) in order to test discrimination of the overlapping routes. In both experiments subjects successfully learned to discriminate between the overlapping routes as evidenced by a higher percentage of target responses than competitor responses (Experiment 1: t_19_ = 8.59, p = 0.00000006; Experiment 2: t_20_ = 6.44, p = 0.000003). In Experiment 1, subjects were more likely to select the competitor destination than one of the ‘other’ destinations (t_19_ = 11.52, p < 0.00000001), indicating that route overlap contributed to memory interference. Error bars reflect +/- SEM. *** p < 0.001

**Figure S2.**
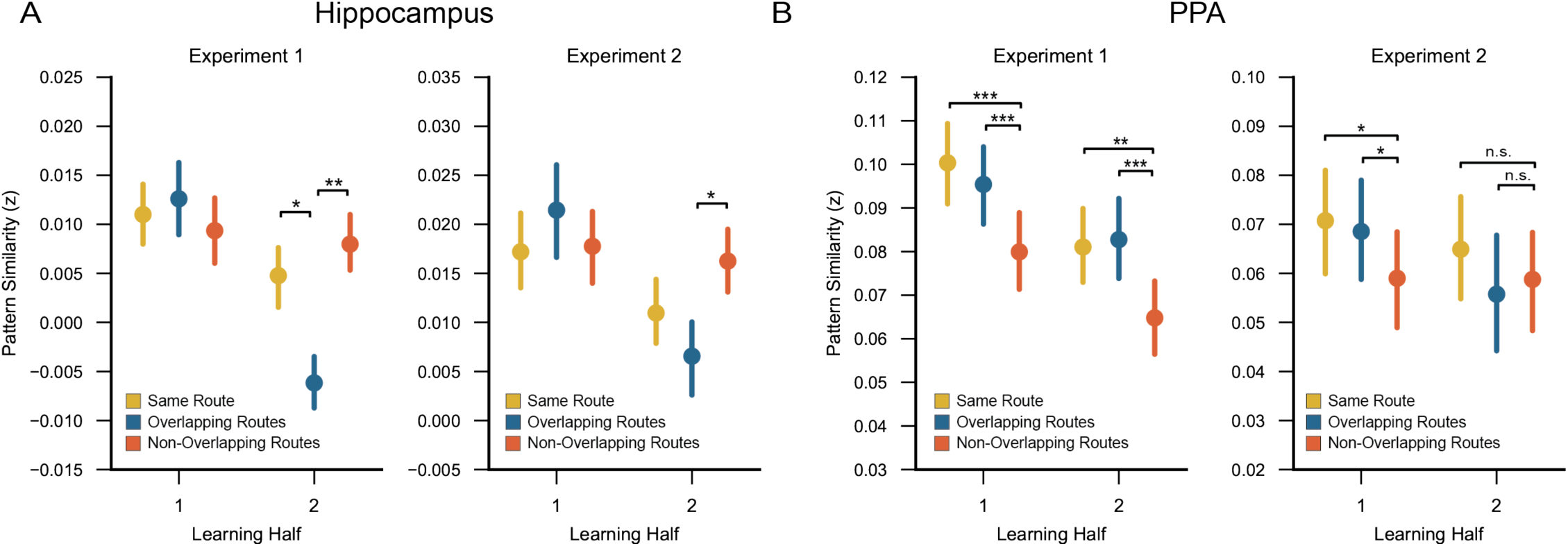
Related to *Figure 3*. Learning-related changes in spatiotemporal pattern similarity (Segment 1 only) for each fMRI Experiment. (A) In each fMRI Experiment, there was a significant learning-related decrease in the similarity of hippocampal representations of overlapping routes relative to non-overlapping routes (Experiment 1: F_1,19_ = 5.99, p = 0.024; Experiment 2: F_1,20_ = 8.02, p = 0.010). Furthermore, the reversal effect (overlapping route similarity < non-overlapping route similarity) was significant in the 2nd half of learning for each Experiment (Experiment 1: t_19_ = 3.03, p = 0.007; Experiment 2: t_20_ = 2.28, p = 0.034). (B) Within PPA, the interaction between learning half (1st vs. 2nd) and overlap (overlapping vs. non-overlapping routes) was not significant in Experiment 1 (F_1,19_ = 0.45, p = 0.51). In both the 1st and 2nd halves of learning, overlapping route similarity was significantly greater than non-overlapping route similarity (1st-half: t_19_ = 4.56, p = 0.0002; 2nd-half: t_19_ = 4.76, p = 0.0001). In Experiment 2, however, the interaction between learning half (1st vs. 2nd) and overlap (overlapping vs. non-overlapping routes) was significant (F_1,20_ = 5.19, p = 0.034), reflecting a relative decrease in overlapping route similarity across learning. Whereas overlapping route similarity was greater than non-overlapping route similarity in the 1^st^ half of learning (t_20_ = 2.27, p = 0.034), there was no difference between overlapping and non-overlapping route similarity in the 2nd half of learning (t_20_ = 0.55, p = 0.59). Error bars reflect +/- SEM. * p < 0.05, ** p < 0.01, *** p < 0.001.

**Figure S3.**
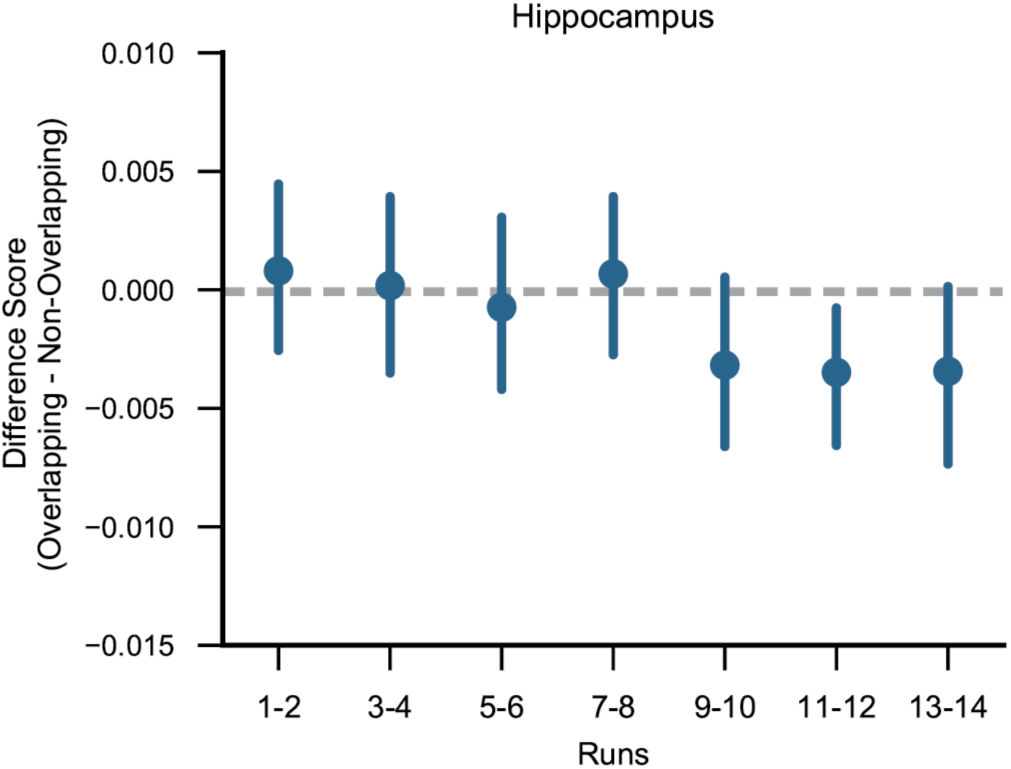
Related to *Figure 3*. Hippocampal spatiotemporal pattern similarity (Segment 1 only) computed every two runs. Qualitatively, there was no evidence for a reversal effect (overlapping route similarity < non-overlapping route similarity) until run 9. However, because each run contained only two repetitions of each route, this analysis was under-powered relative to the main analyses split by learning half.

**Figure S4.**
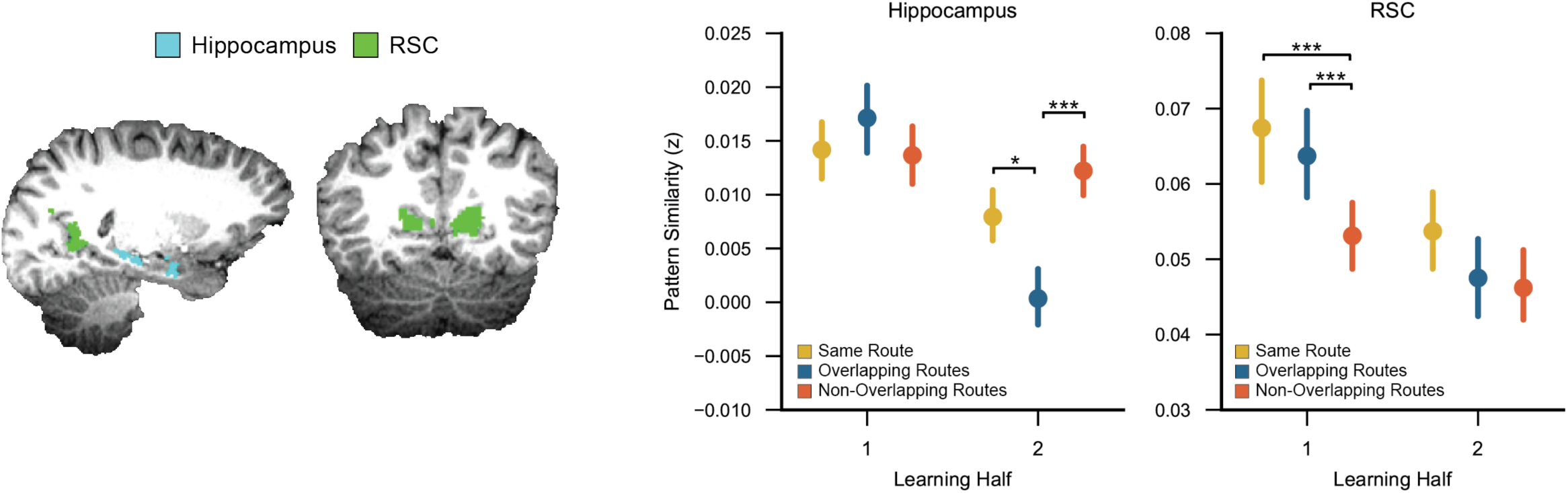
Related to *Figure 3*. Comparison of learning-related changes in spatiotemporal pattern similarity (Segment 1 only) for hippocampus vs. retrosplenial cortex (RSC). In addition to PPA (our primary control region), retrosplenial cortex (RSC) has also been implicated in scene processing, spatial navigation, and episodic memory retrieval (Vann et al., 2009; Epstein, 2008; Marchette et al., 2014). Within RSC, overlapping route similarity decreased across learning, relative to non-overlapping route similarity (F_1,39_ = 4.45, p = 0.041). In the 1st half of learning, overlapping route similarly was significantly greater than non-overlapping route similarity (F_1,39_ = 16.10, p = 0.00026). In the 2nd half, however, there was no difference between overlapping and non-overlapping route similarity (F_1,39_ = 0.11, p = 0.74). Thus, although overlapping route similarity decreased relative to non-overlapping route similarity, there was no evidence of the reversal effect (overlapping route similarity < non-overlapping route similarity) that was observed in the hippocampus. There was also no learning-related changes in overlapping route similarity relative to same route similarity (F_1,39_ = 0.18, p = 0.67). Overlapping route similarity did not differ from same route similarity in either the 1st half (F_1,39_ = 1.41, p = 0.24) or 2nd half of learning (F_1,39_ = 1.91, p = 0.18). For comparison, data from the hippocampus are shown here (identical to Figure 3C). The difference in representational structure across hippocampus and RSC was confirmed by a significant interaction between overlap (overlapping vs. non-overlapping) and region (hippocampus vs. RSC) within the 2nd half of learning (F_1,39_ = 10.23, p = 0.0027). Thus, although RSC patterns did change with learning, the representational end-states of learning were qualitatively different across hippocampus and RSC. Error bars reflect +/- SEM. * p < 0.05, ** p < 0.01, *** p < 0.001

**Figure S5.**
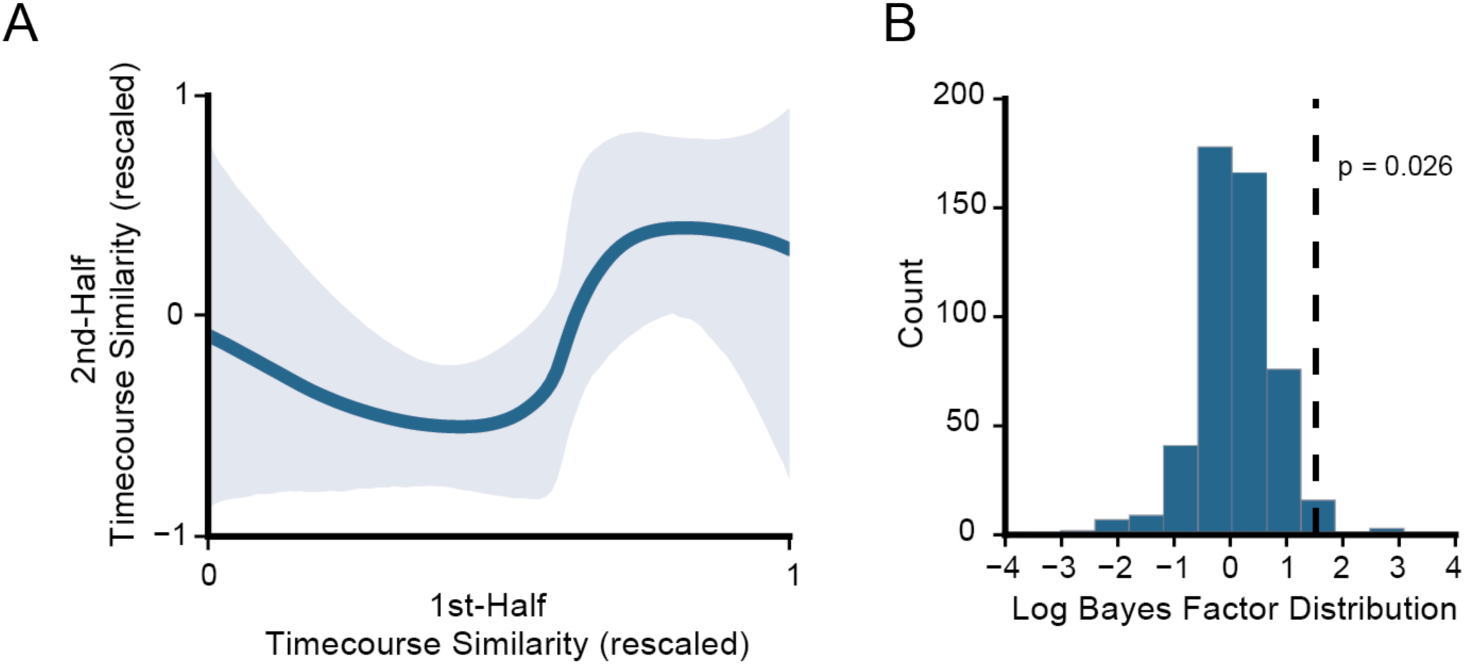
Related to *Figure 5*. Bayesian curve-fitting analysis. To more formally assess the non-monotonic relationship between 1st-half and 2nd-half timecourse similarity, we used a Bayesian curve-fitting algorithm–the Probablistic Curve Induction and Testing Tooblbox (P-CIT) (Detre et al., 2013) that was specifically developed to test for non-monotonic plasticity. Relative to quadratic trend analyses, the P-CIT algorithm allows for a more detailed specification of a predicted curve shape, by explicitly including a set of curve parameters. In our case, the parameters describe the relationship between first-half timecourse similarity (x-axis) and second-half timecourse similarity (y-axis). We parameterized the predicted curve shape using previously described parameters that reflect the prediction of non-monotonic plasticity (Kim et al., 2014). Specifically, the predicted curve was defined as one in which the function, when moving from left to right, drops below the initial start value and then rises above the start value. The first step of the P-CIT algorithm is to estimate a curve shape given the data. To accomplish this, the algorithm estimates a probability distribution over possible curves, conditional on the observed data, by randomly sampling curve shapes and then assigning each sampled curve an importance weight indicating how well the curve’s shape fit the observed data. It then estimates a curve by averaging the sampled curves together, weighted by their importance values. The next goal of the algorithm is to evaluate the level of evidence in favor of the predicted curve shape. It does so by labeling each sample curve as theory consistent (in our case, if it drops below the starting value and then rises above the starting value) or inconsistent, and then computes a log Bayes factor value that represents the log ratio of evidence in favor of or against the predicted shape (Lewis-Peacock et al., 2014). Positive log Bayes factor values indicate greater evidence in favor of the theory. For this analysis, we re-binned all of the 1st-half timecourse similarity values into 60 bins (5 voxels per bin) in order to allow for greater variability in the observed curve shape. This analysis used data aggregated across all subjects. (A) The estimated curve was consistent with the predicted curve shape (log Bayes factor = 1.51) and explained a significant amount of variance in the actual (*X*_2_ = 11.13, p = 0.0008). Shaded area reflects the 90% credible interval. (B) We next ran a permutation test to estimate the null distribution of log Bayes Factor values. Out of 500 permutations, only 2.6% yielded log Bayes factor values that matched or exceeded the value obtained from the un-permuted data, indicating that it was unlikely to obtain this level of support for the predicted curve shape by chance. Finally, to assess the population-level reliability of the non-monotonic curve we ran a bootstrap resampling test in which we iteratively resampled data from subjects with replacement and then computed the log Bayes factor value for each iteration. Four-hundred and eighty-eight of the 500 bootstrap iterations (97.6%) yielded positive log Bayes factor values. Thus, the curve-fitting analyses provided additional evidence for a nonmonotonic relationship between voxel overlap at the beginning vs. end of learning: that is, hippocampal voxels that were ‘moderately shared’ across overlapping routes at the beginning of learning were the ‘least shared’ by the end of learning.

